# 16S rRNA Gene Amplicon Profiling of the New Zealand parasitic blowfly *Calliphora vicina*

**DOI:** 10.1101/2021.03.12.435192

**Authors:** Nikola Palevich, Paul H. Maclean, Luis Carvalho, Ruy Jauregui

## Abstract

Here, we present a 16S rRNA gene amplicon sequence data set and profiles demonstrating the bacterial diversity of larval and adult *Calliphora vicina*, collected from Ashhurst, New Zealand (May 2020). The three dominant genera among adult male and female *C. vicina* were *Serratia* and *Morganella* (phylum *Proteobacteria*) and *Carnobacterium* (phylum *Firmicutes*), while the larvae were also dominated by the genera *Lactobacillus* (phylum *Firmicutes*).

## ANNOUNCEMENT

Ectoparasitic flies (blowflies) are a significant animal welfare and production issue for farmers worldwide (1). Control of blowflies is problematic because the flies are unpredictable and highly mobile, and strike (or myiasis) is difficult to see initially but has an immediate impact on animal production and welfare. Currently control relies heavily on the prophylactic application of long-acting chemicals to all sheep, but this approach is increasingly under threat due to development of resistance to current treatments (2, 3). *Calliphora vicina* NZ_CalVic_NP (4, 5) was selected for microbiome assessment as a representative of a New Zealand field strain of *C. vicina*. In this study, we have investigated the larval, adult male and adult female bacterial microbial profiles of *C. vicina* to gain a better understanding of the microbial communities of blowflies targeting the development of new interventions such as probiotics, bioactive compounds, vaccines or insecticides.

The *C. vicina* specimen larvae were collected from a farm site in Ashhurst area in New Zealand (40°18′ S, 175°45′ E). Lab reared blowflies were maintained on beef liver as protein source and a 10% sugar solution, with the procedures for blowfly propagation and sample preparation were based done according to Dear J.P. (1985). To remove surface adherent bacteria from lab reared *C. vicina*, pools of larvae, entire adult male and adult female were separated and washed twice in sterile phosphate-buffered saline (PBS, pH 7.4), snap frozen in liquid nitrogen, and transferred to −80 °C storage prior to DNA extraction. Genomic DNA for metagenomic 16S rRNA gene amplicon sequencing of the V3-V4 hypervariable region was isolated from *C. vicina* pooled samples of 100 larvae as well as 10 entire adult males and females per replicate (*n*=5 for each). High molecular weight genomic DNA was prepared using a modified phenol:chloroform protocol recently applied to difficult samples such as parasitic roundworms (7, 8), fastidious anaerobic rumen bacteria (9–11) and spore-forming psychrotolerant *Clostridium* isolated from spoiled meat (12, 13). A DNA library was prepared using the Illumina 16S V3-V4 rRNA library preparation method (Illumina, Inc., San Diego, CA) according to the manufacturer’s instructions (20), and sequenced on the Illumina MiSeq platform with the 2× 250 bp paired-end (PE) reagent kit v2 producing a total of 3,017,007 PE raw reads.

The processing of the amplicon reads followed a modified form of the pipeline described in (21). The reads produced by the sequencing instrument were paired using the program FLASH2 (22). Paired reads were then quality trimmed using Trimmomatic 0.38 (23). The trimmed reads were reformatted as fasta, and the read headers were modified to include the sample name. All reads were compiled in a single file, and the Mothur (24) program suite was used to remove reads with homopolymers longer than 10 nt and to collapse the reads into unique representatives. The collapsed reads were clustered using the program Swarm (25). The clustered reads were filtered based on their abundance, keeping representatives that were a) present in one sample with a relative abundance >0.1%, b) present in >2% of the samples with a relative abundance >0.01% or c) present in 5% of the samples at any abundance level. The selected representatives were annotated using Qiime (26) with the Silva database v138 (27). The annotated tables were then used for downstream statistical analysis. The predominant phyla in all samples were Proteobacteria (Fig. 1) and at the genus level, *Serratia*, *Morganella* and *Carnobacterium*, while the larvae were also dominated by *Lactobacillus* (phylum Firmicutes).

**FIG 1.**
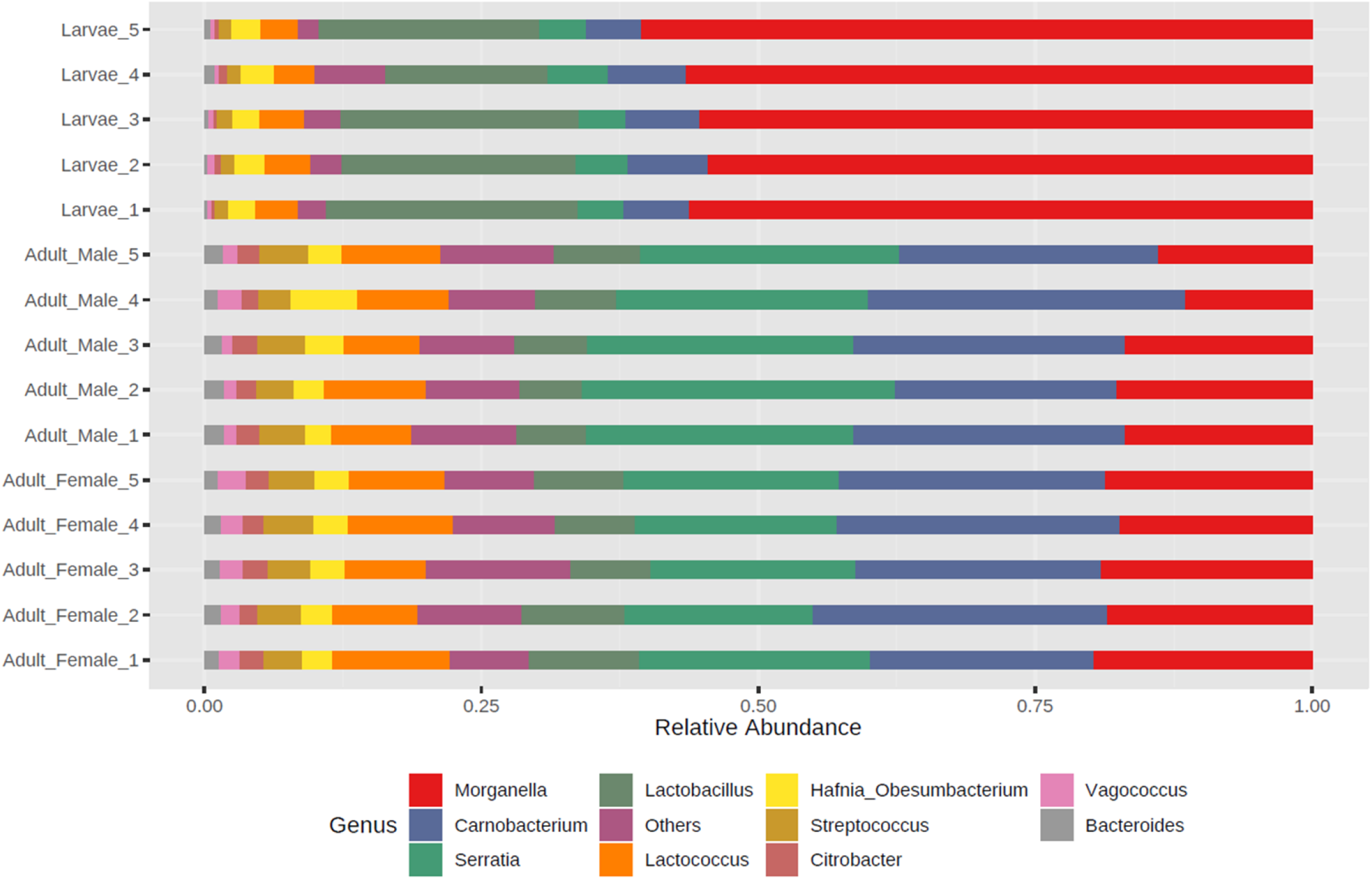
The taxonomic composition of the dominant bacteria of New Zealand *C. vicina*. Relative abundance of the dominant bacterial genera obtained from 16S rRNA sequencing of *C. vicina* field strain NZ_CalVic_NP larvae, adult males and female samples. Genera with a relative abundance of less than 1% and unassigned amplicon sequence variants were grouped together as Others.

The metagenomic 16S rRNA gene amplicon sequencing of *C. vicina* field strain NZ_CalVic_NP reported here is a valuable resource for future studies investigating the bacterial genetic mechanisms associated with flystrike. Management of flystrike in a world increasingly demanding fewer inputs of synthetic chemicals to food producing animals will be challenging. Equally, this research is important owing to the diminished efficacy demonstrated by current blowfly treatments due to the emergence of resistance in blowflies against many classes of insecticides.

## Data availability

The 16S rRNA gene amplicon sequence data have been deposited in the GenBank Sequence Read Archive (SRA) under the BioProject accession number PRJNA667961.

## ACKNOWLEDGEMENTS

We thank Xiaoxiao Lin for assistance with the DNA sequencing and Paul Candy for sample collection. This research was supported by the Agricultural and Marketing Research and Development Trust (AGMARDT) Postdoctoral Fellowships Programme [No. P17001] and the AgResearch Ltd Strategic Science Investment Fund (SSIF) [No. PRJ0098715] of New Zealand.

